# Metabolic niches: MALDI imaging reveals lipidomic heterogeneity in the human heart

**DOI:** 10.64898/2026.06.18.732349

**Authors:** Kayla M. Osumi, Evan P. Kransdorf, Anja Karlstaedt, Elizabeth K. Neumann

## Abstract

The human heart is characterized by aberrant lipid accumulation and remodeling during periods of stress and disease, yet spatially resolved lipidomic profiling of the human heart remains unreported. Here, we employ matrix-assisted laser desorption/ionization mass spectrometry imaging (MALDI-MSI) to map lipid distributions across five anatomical regions of healthy human donor hearts (left and right atrium, left and right ventricle, and interventricular septum). Cryosections of donor hearts from healthy subjects were thaw-mounted onto indium tin oxide slides, coated with 2,5-dihydroxyacetophenone, and analyzed on a Bruker timsTOF fleX mass spectrometer in positive and negative ionization modes (50–1850 *m/z*; 20 μm raster). Serial sections were stained with hematoxylin and eosin to enable histological co-registration with lipid distributions. We identified 150 unique lipid species with acyl-chain resolution across four glycerophospholipid classes — phosphatidylcholine (PC), phosphatidylglycerol (PG), phosphatidylethanolamine (PE), and phosphatidylserine (PS) — with abundances and spatial distributions differing significantly between regions. H&E co-registration further resolved epicardial and myocardial compartments, enabling tissue-specific lipid mapping. These findings establish the first spatially resolved lipidomic atlas of the human heart and provide a framework for identifying region-specific lipid biomarkers of cardiovascular disease.

## INTRODUCTION

Cardiovascular disease (CVD), including heart failure, is the most common cause of death in the United States.^1^ Lipids play a critical role in energy provision, and as essential structural components of cellular membranes that regulate ion transport, signal transduction, and membrane integrity. The development of CVD is characterized by systemic and cardiac-specific lipid alterations, but prior studies have focused mainly on storage lipids (e.g., diacylglycerides and triacylglycerides). The spatial organization of membrane lipids — including phospholipids and sphingolipids, key determinants of cellular structure and signaling — remains poorly defined in the human heart.

Early subcellular lipidomic studies revealed the predominance of choline and ethanolamine plasmalogens in membrane isolations from the canine heart ^2^. Complementary analyses of human cardiac biopsy specimens^3,4^ established that the major phospholipid classes of the human heart include phosphatidylcholine (PC), phosphatidylethanolamine (PE), diphosphatidylglycerol, and sphingomyelin, with sex and age having minimal influence on the relative distribution of these classes. However, these early studies did not fully resolve all lipid classes and lacked insights into the acyl-chain compositions.

Phospholipids are not uniformly distributed across subcellular compartments, and perturbations in this composition are closely linked to cardiac dysfunction^5–7^. Further, phospholipid classes are asymmetrically distributed across sarcolemmal and mitochondrial membrane, with negatively charged phosphatidylserine and phosphatidylinositol localized exclusively to the inner cytoplasmic leaflet, and sphingomyelin predominantly at the outer leaflet. This asymmetry has functional consequences for ion exchanges, energy provision and nutrient exchange across organelles. Changes in these compositions have been associated with increased cardiac injury in response to ischemia-reperfusion^8^. The spatial organization of phospholipids is a critical determinant of cardiac function.

Recent advances in mass spectrometry-based lipidomics has expanded the ability to characterize and quantify the lipidome in mammalian tissue. Untargeted liquid chromatography and mass spectrometry (LC-MS)-based lipidomics of human cardiac tissue has revealed distinct lipid profiles between healthy and ischemic subjects, with PE among the most significantly downregulated species in the ischemic heart, pointing to PE remodeling as a hallmark of pathological cardiac adaptation^9^. Likewise, analysis of human cardiac tissue explants revealed lower total linoleic acid content alongside elevated polyunsaturated fatty acid species, suggesting increased delta-6-desaturase activity as a feature of disease-associated phospholipid remodeling. Collectively, these studies establish that phospholipid composition changes are a hallmark of cardiac remodeling, yet these studies are largely limited to bulk tissue analysis that sacrifice spatial information.

Despite these advances, the spatial distribution of phospholipid species across anatomical regions of the human heart remains uncharacterized. Although it is well established that cardiac tissue is enriched in plasmalogens to a considerably greater extent than other species^10^, no lipidomic studies have thoroughly evaluated the lipidomes of individual cardiac chambers, despite each chamber having tremendously different roles in cardiac physiology. Furthermore, tissue-specific lipidomic fingerprints have been demonstrated across multiple organs, with glycerophospholipids constituting the most abundant lipid class and displaying tissue-specific distributions at the molecular species level^5,11^. However, equivalent region-resolved data for the human heart are lacking. Matrix-assisted laser desorption/ionization mass spectrometry imaging (MALDI-MSI) addresses this gap by generating spatially preserved mass spectra across tissue sections at defined raster increments, enabling direct mapping of lipid distributions onto tissue architecture without the need for extraction or homogenization^12^. MALDI-MSI has been established as a powerful analytical tool for research, capable of resolving spatial distributions of lipids within tissue sections and has been applied to characterize lipid alterations in preclinical animal models of myocardial infarction and ischemia-reperfusion injury^13^. However, its application to the healthy human heart across multiple anatomical regions has not been reported.

Here, we employ MALDI-MSI to generate the first spatially resolved lipidomic atlas of the healthy human heart, profiling phospholipid distributions across five anatomical regions (e.g. left and right atria, left and right ventricles, and interventricular septum) in positive and negative ionization modes. By integrating spatial lipid mapping with histological co-registration, we resolve region- and tissue-specific phospholipid distributions, providing a reference framework for future studies of lipid dysregulation in cardiac disease states.

## METHODS

### Human heart tissue preparation

Adult human hearts were procured as donor human hearts from UNOS Transplant Services between May and August 2024. Experimental protocols were approved by the Smidt Heart Institute Tissue Repository (Protocols: CS-IRB Pro00010979 and Pro00011910). Informed consent was obtained from all patients. Human heart tissue samples were obtained from donor patients who were rejected for transplantation due to organ size, lack of an organ recipient, or ischemia time. Explanted hearts were cardioplegically arrested with a high-potassium solution (in mmol/L: NaCl 110, CaCl_2_ 1.2, KCl 16, MgCl_2_ 16, NaHCO_3_ 10; Sigma-Aldrich, St. Louis, MO, USA) and cooled to 4 °C in the operating room following aortic cross-clamp. Myocardium from five anatomical heart regions (left and right ventricle, left and right atrium, and intraventricular septum) was dissected and frozen over dry ice before being embedded in a carboxymethylcellulose solution. Ischemia time between heart clamping and tissue processing for MALDI-MSI was less than 3.5 hours.

### MALDI MSI Sample Preparation and Analysis

Chemicals for sample processing were purchased from MilliporeSigma without further purification unless otherwise specified. Tissue was cryosectioned to a thickness of 12 μm at −20 °C (Leica Biosciences, Wetzlar, Germany) and thaw-mounted onto a conductive indium tin oxide slide (Delta Technologies, Limited, Loveland, Colorado). The slide was coated with a 10 mg/mL 2,5-dihydroxyacetophenone (DHA) in 70% acetonitrile using an M3+ TM sprayer (HTX Technologies, LLC, Chapel Hill, NC). Sprayer parameters were 8 passes at 120 μL/min, 1200 mm/min velocity, 2.5 mm track spacing, 15 psi pressure, and 50 °C. MALDI MSI experiments were carried out using a timsTOF fleX MALDI mass spectrometer (Bruker Scientific, Billerica, MA) in positive and negative polarity with a 20 μm raster size. Positive mode parameters included 250 shots per pixel, 87% laser energy (0% global attenuator), and a mass range of 50 to 1850. Negative mode parameters included 100 shots per pixel, 66.9% laser energy (0% global attenuator), and a mass range of 50 to 3000. Beam scan for both modes set to 6 μm x 6 μm. Ion images and heart region-specific average spectra were generated using SciLS (Version 2026a Pro), Bruker Scientific, Billerica, MA). Internal calibration for spectra was conducted on *m/z* lists generated before putative identifications using the LipidMAPS database^14,15^. Raw data from timsTOF fleX runs were converted to an imzML file type using TIMSCONVERT (v2.0.0)^16^. Converted data was processed using the Cardinal package (v3.8.3) in R (v4.4.3).

### Histology and Hematoxylin and Eosin (H&E) staining

Tissue sections for histology were prepared from frozen heart tissue sections for MALDI-IMS imaging. Frozen tissue blocks were sectioned at 12 μm thickness using a microtome and mounted onto glass slides. Sections were incubated in isopropanol for 1 min to remove residual DHA matrix, air-dried, and stained with hematoxylin followed by eosin according to standard protocols. After staining, the slides were dehydrated, cleared, and coverslipped using glycerol wet-mount medium (Cat. No. RS-STB-097, Rs’ Science). Images were acquired using an Axio Scan.Z1 slide scanner (Zeiss, Oberkochen, Germany) at 20x magnification under identical imaging settings across samples. Representative images are shown in the figures.

### Uniform Manifold Approximation and Projection (UMAP) Analysis

Uniform Manifold Approximation and Projection (UMAP) was used for dimensionality reduction and visualization of sample clustering. Principal component analysis (PCA) was first conducted, and the first 100 principal components were retained as input for UMAP to reduce noise and computational complexity while preserving the major sources of variation in the dataset. UMAP was conducted using uwot package (v0.2.3)^17^ in R (v4.4.3). The low-dimensional embedding was generated using the following parameters: number of nearest neighbors = 15, minimum distance = 0.01, distance metric = Euclidean, and K-means clustering = 8. A two-dimensional embedding was computed for visualization purposes. Samples were colored by experimental group, and clustering patterns were assessed qualitatively by the separation of groups in the two-dimensional UMAP space.

### UpSet plot

The UpSet plot was generated using the R package UpSetR^18^. To identify discriminating *m/z* values per cluster, we used a log2 fold change to compare the average spectra of pixels associated with one cluster against those of another. The top five discrimination m/z values per cluster were obtained after filtering by mass defect of 0.4 – 0.6 to capture lipid biology.

### Statistical Analysis

Partial Least Squares Discriminant Analysis (PLS-DA) was conducted on normalized metabolomics data using the R package mdatools (v0.14.2). The following R packages were used for general data analysis: base (v4.3.2), dplyr (v1.1.4), readxl (v1.4.3), tibble (v3.2.1), tidyverse (v2.0.0); and Data Visualization: ggplot2 (v3.5.2) and pheatmap (v1.0.13). Claude (Anthropic) was used to assist with R code development.

### Data Availability

Visualization of the entire dataset are available through a shinyapp: *https://karlstaedtlab.shinyapps.io/MALDI-Imaging-Human-Heart/*

## RESULTS

### Regional Lipidomic Profiling of the Human Heart by MALDI-MSI

We conducted MALDI-MSI on tissue sections from five anatomical regions of four healthy donor hearts, generating spatially preserved lipid profiles in both positive and negative ionization modes. Tissue samples were obtained from four human heart donors (ejection fraction >65%) and embedded in MALDI compatible matrix (see Methods for details, (**Figure 1A**). In positive mode, the number of putatively identified lipid species per region was as follows: 85 in the right atrium (RA), 75 in the left atrium (LA), 99 in the right ventricle (RV), 78 in the left ventricle (LV), and 90 in the interventricular septum (IVS). In negative mode, 66 lipids were identified in the RA, 73 in the LA, 78 in the RV, 74 in the LV, and 66 in the IVS (**Figure 1B**). Detected lipid species spanned multiple classes, including glycerophospholipids (PC, PE, PS, PI, PG, PA), sphingolipids (SM, CerPE, CerP, HexCer, SHexCer), storage glycerolipids (TG, DG), and plasmalogens (PC, PE, PA, PG, PI, and PS ether-linked species).

**Figure 1.**
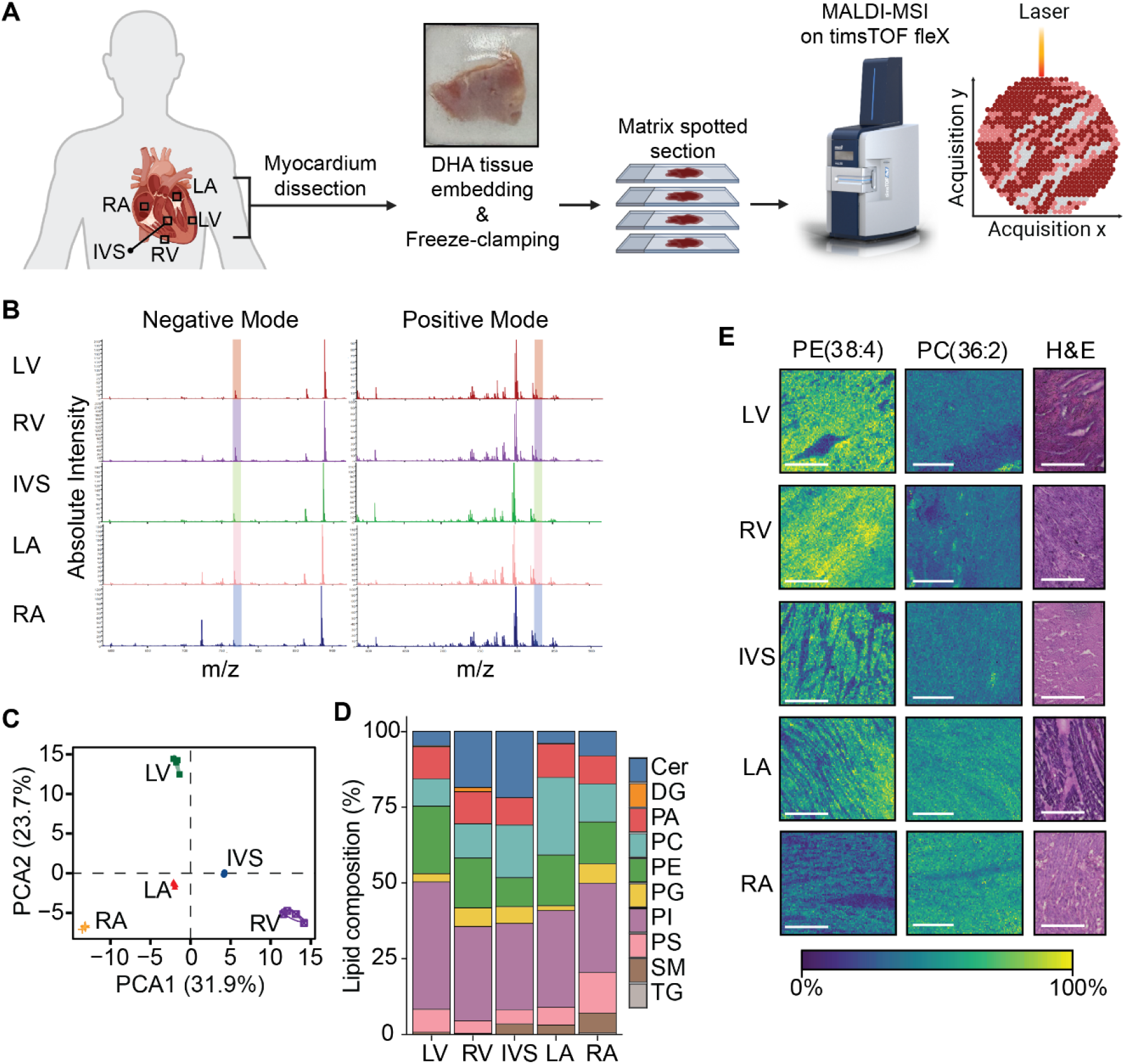
MALDI-IMS workflow to reveal lipidomic landscape of the human heart. (**A**) Schematic of tissue acquisition and MALDI-IMS workflow. (**B**) Representative negative and positive average spectra for cardiac regions. (**C**) Principal component analysis of lipid abundances based on MALDI-IMS across anatomical heart regions. (**D**) Lipid composition analysis showing the relative distribution (%) of the lipid classes across the five anatomical heart regions (LV, RV, RA, LA, and IVS). (**E**) Representative H&E’s taken of the five anatomical regions from a subsequent tissue section. Abbreviations: Cer, ceramide; DG, diacylglycerols; IVS, intraventricular septum; LA, left atrium; LV, left ventricle; PA, phosphatidic acid; PI, phosphatidylinositol; PC, phosphatidylcholine; PS, phosphatidylserine; PE, phosphatidylethanolamine; RA, right atrium; RV, right ventricle; SM, sphingomyelin; TG, triacylglycerols.

Principal component analysis (PCA) demonstrated clear separation between the five anatomical regions (**Figure 1C**), with the primary sources of variation attributable to glycerophospholipid classes — including PA, PI, PC, PS, and PE — as well as sphingolipids including ceramide (Cer) and sphingomyelin (SM) (**Figure 1D**). Sphingomyelins were markedly enriched in the atria (LA and RA) relative to the ventricles, where they were virtually absent. Diacylglycerols (DG) and triacylglycerols (TG) collectively accounted for less than 5% of detected lipids in the LV and RV, consistent with the predominance of membrane phospholipids over storage lipids in healthy ventricular myocardium.

Ion images revealed spatially heterogeneous lipid distributions that differed both between and within anatomical regions (**Figure 1E**). For example, SM(34:1;O2) displayed a discrete, localized distribution in the RA, LA, LV, and IVS, while in the RV the same species was dispersed throughout the tissue, suggesting region-specific functional roles. Comparison of average mass spectra across regions further highlighted quantitative differences in lipid abundance (**Figure 1E**). The mean pixel intensity of PE(38:2) in positive mode differed substantially across regions: RV 50.87 ± 12.53, LV 29.79 ± 12.73, LA 29.21 ± 11.83, IVS 22.70 ± 15.03, and RA 16.71 ± 1.62. The RV exhibited the highest PE(38:2) abundance while the RA showed markedly lower intensity relative to all other regions. In negative mode, PC(36:2) abundance also varied significantly across regions: LA 31.38 ± 12.04, RA 23.85 ± 7.04, RV 18.65 ± 2.56, LV 15.36 ± 1.68, and IVS 14.94 ± 2.59. The LA was most enriched in PC(36:2), while the IVS showed the lowest abundance, with ion images corroborating these quantitative differences through visual comparison of signal intensity across regions.

### Shared and Unique Lipid Species Across Regions

UpSet plot analysis of putative lipid assignments quantified the degree of overlap and uniqueness between regions (**Figure 2**). In negative mode (**Figure 2A**), 17 lipid species were detected across all five regions, while 20 were unique to the RV, 20 to the RA, 16 to the IVS, 13 to the LV, and 13 to the LA. In positive mode (**Figure 2A**), no lipids were shared across all five regions, reflecting greater inter-regional heterogeneity compared to negative mode. Region-unique species were most abundant in the ventricles in positive mode, with 37 unique to the RV and 40 to the LV.

**Figure 2.**
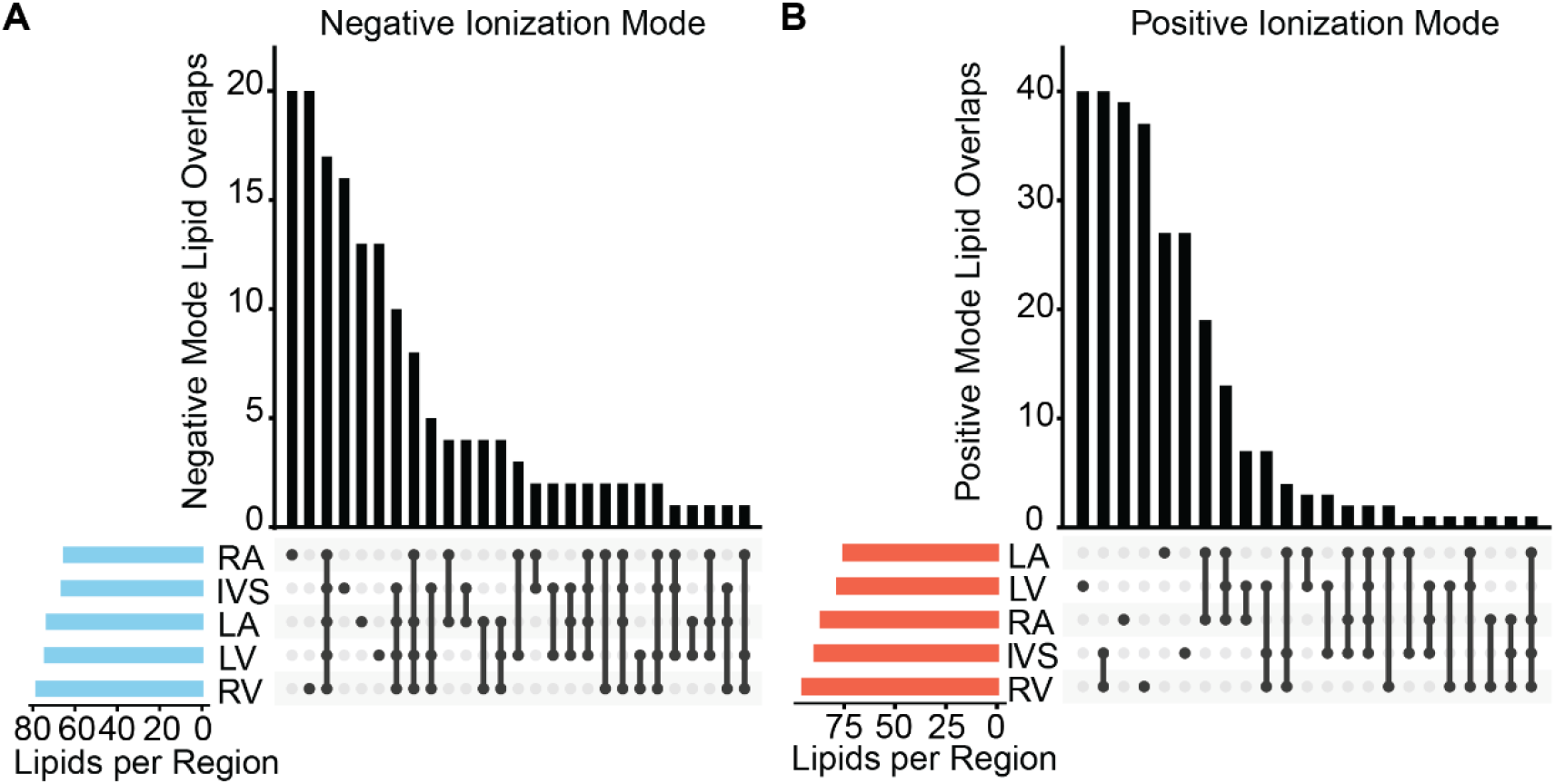
Landscape of Lipid Detection Across Anatomical Regions. (**A**–**B**) UpSet plots illustrating the intersecting distributions of putatively assigned lipids detected in negative ionization mode (**A**) and positive ionization mode (**B**). Horizontal bar graphs (left) denote the total lipidome size per anatomical region. Vertical bar graphs (right) represent intersection sizes, reflecting the number of lipids exclusively detected within a single region or co-detected across multiple regions. Filled circles beneath each intersection bar indicate the contributing regions, with vertical connectors denoting shared lipid species between two or more regions. n = 4 healthy donor hearts per region.

### UMAP Clustering Identifies Spatially Reproducible Lipid Communities

Phospholipid synthesis is closely linked across species allowing remodeling and acyl-chain modifications (**Figure 3A**). To further assess spatially resolved lipid features, we applied Uniform manifold approximation and projection (UMAP) to negative mode data. Our analysis revealed partial overlap between the five anatomical regions (**Figure 3B**), suggesting a broadly shared phospholipid framework across the heart. k-means clustering on UMAP embedding resolved eight discrete clusters (**Figure 3B**), which were reproducible across all four donor samples. The top five discriminating lipid species per cluster, determined by Log2 fold change in mean pixel intensity relative to all other clusters, are visualized in a heatmap (**Figure 3C**). Hierarchical clustering of these features revealed cluster-specific enrichment of PE(36:1), PE(38:4), PS(36:1), PI(36:1), and PI(36:4), which are primarily comprised of FA18:0, FA18:1, FA18:2, and FA16:0. These findings suggest that these lipid species allow separation of anatomical regions.

**Figure 3.**
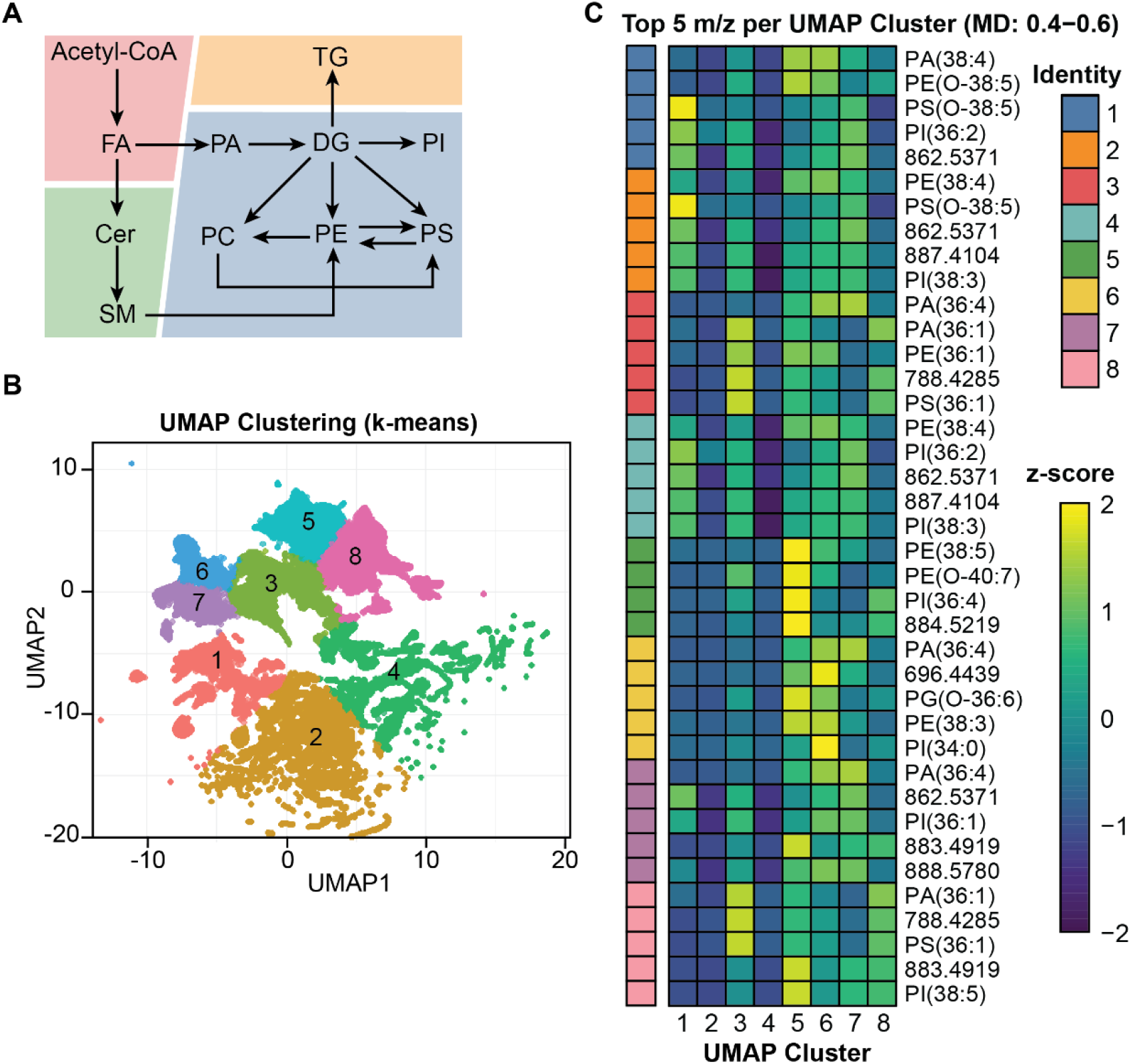
Spatial Mapping of De Novo Phospholipid and Sphingolipid Biosynthesis Pathways. (**A**) Schematic illustration of the de novo biosynthetic pathways for phospholipids and sphingolipids, highlighting key enzymatic steps, metabolic intermediates, and phospholipid remodeling reactions. (**B**) UMAP dimensionality reduction followed by k-means clustering (k = 8) was applied to negative ionization mode mass spectrometry imaging data to spatially resolve lipid heterogeneity across anatomical regions, with each pixel colored by cluster assignment. (**C**) The top five discriminating lipid species per cluster, determined by Log2 fold change in mean pixel intensity relative to all other clusters, were subjected to unsupervised hierarchical clustering and visualized as a heatmap. The relative lipid enrichment between clusters using z-scored cluster mean intensities is shown for all eight UMAP clusters. Color intensity reflects the magnitude and directionality of enrichment, with rows representing lipid features and columns representing UMAP-derived clusters.

### Network analysis reveals lipidomic profiles in the heart

We further inspect the spatial distribution of acyl-chain composition 36:1, 36:4 and 38:4 across anatomical regions. The spatial distribution of the 36:1 acyl composition provided the clearest separation between anatomical regions across all quantified phospholipid and sphingolipid classes (**Figure 4A**). 36:4 and 38:4 were enriched in PA and PI species, respectively across all classes. Thus, these compositions allow lipid class separation rather than anatomical distribution. We utilized network-based analysis of lipid species interactions based on known de novo synthesis and remodeling patterns using LIPIDMAPS database. The scaled fractional enrichments of 36:1, 36:4 and 38:4 species were used to generate spatial networks for anatomical regions (**Figure 4B**). Network analysis revealed regional patterns and links for LV and RV.

**Figure 4.**
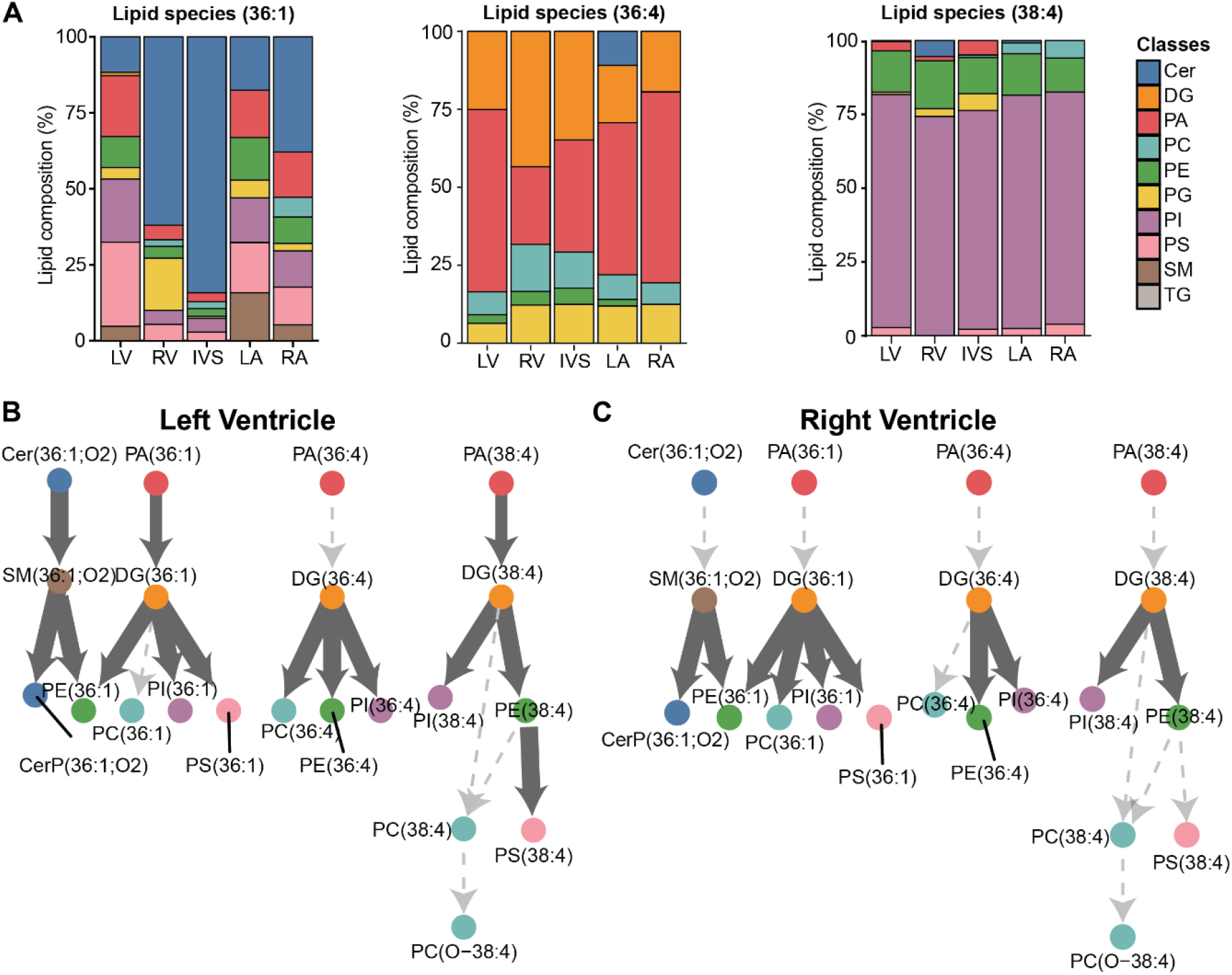
Regional Heterogeneity of Lipid Class Distributions Revealed by Species-Level Network Analysis. (**A**) Lipid composition analysis showing the relative distribution (%) of the 36:1, 36:4, and 38:4 lipid species across the five anatomical heart regions (LV, RV, RA, LA, and IVS). (**B**) Lipid species interaction network illustrating the fractional contribution of individual lipid species within each lipid class across LV and RV. Networks were constructed using lipid class annotations from the LIPID MAPS structure database. Nodes represent individual lipid species and are colored by lipid class. Edges represent the fractional abundance of each species relative to its parent lipid class, with edge thickness scaled to fractional contribution binned across quartile ranges, reflecting increasing relative abundance. Edges with a fractional contribution of zero are rendered as dashed lines with reduced opacity to denote trace or absent species. Networks were independently constructed for each anatomical heart region to capture region-specific lipid class compositions.

## DISCUSSION

The present study provides the first spatially resolved lipidomic profiling of the healthy human heart across five anatomical regions using MALDI-MSI, revealing region-specific differences in phospholipid abundance, class composition, and spatial distribution. While previous lipidomic studies of the heart have primarily employed bulk LC-MS approaches that yield molecular detail without spatial context, MALDI-MSI uniquely preserves the spatial coordinates of each acquired spectrum, enabling direct mapping of lipid identity and abundance onto tissue architecture^13,14^.

PCA-driven separation between anatomical regions indicates that the lipidomic identity of each cardiac region is distinct, driven primarily by differences in glycerophospholipid and sphingolipid composition. The pronounced enrichment of sphingomyelins in the atria relative to the ventricles is consistent with previously reported differences in membrane composition between chamber types and may reflect differences in electrophysiological properties and signaling requirements between these functionally distinct compartments. The low abundance of storage glycerolipids (TG and DG) in the ventricles corroborates the well-established metabolic phenotype of the healthy ventricular myocardium, which relies predominantly on fatty acid oxidation rather than lipid storage, and in which excessive TG accumulation is a known feature of pathological remodeling.

The regional differences in PE(38:2) abundance are particularly notable. PE constitutes approximately 30% of the mitochondrial membrane and plays a central role in maintaining mitochondrial ultrastructure and respiratory chain function^19^. Mitochondria are more abundant in ventricular than atrial myocardium^20^, consistent with the greater energy demands of ventricular contraction — a relationship that may underlie the markedly higher PE(38:2) intensity observed in the RV compared to the RA. Similarly, the enrichment of PC(36:2) in the LA relative to other regions, and its relatively low abundance in the IVS, may reflect region-specific differences in membrane remodeling associated with the distinct hemodynamic environments of these chambers. Together, these quantitative differences indicate that lipid abundance, not merely lipid identity, is differentially regulated across anatomical regions in a manner that likely reflects underlying functional specialization.

Notable, we did not observe lipid species that are shared across all five regions in positive mode, which contrasts with the 17 shared species identified in negative mode. These findings suggest that the positive mode lipidome, which is enriched in PC and SM species, exhibits greater inter-regional heterogeneity than the negative mode lipidome. This may reflect the greater structural and signaling diversity of choline-containing phospholipids and sphingomyelins across cardiac compartments. The reproducibility of k-means clusters across all four donors further supports the conclusion that the spatially defined lipid communities identified here reflect genuine biological organization rather than technical variation. The cluster-specific enrichment of PE, PS, and PI species bearing the 36:1 and 38:4 acyl compositions suggests region-specific activation of de novo phospholipid biosynthetic pathways, with PA serving as the common precursor for the divergent synthesis of PE, PS, and PI through the CDP-diacylglycerol and Kennedy pathways.

Several limitations of this study merit consideration. Putative lipid identifications were assigned based on accurate mass alone without tandem MS confirmation, and assignments should be interpreted as putative pending validation by MS/MS fragmentation. The study was conducted on healthy donor hearts and does not capture disease-associated lipidomic remodeling. Future studies should extend this spatially resolved framework to pathological cardiac states (e.g. heart failure, ischemic cardiomyopathy, and atrial fibrillation) to identify region-specific lipid biomarkers with diagnostic or mechanistic relevance. Our study revealed region-specific lipidomic signatures in the healthy human heart, providing potential biomarkers and establishing a resource for the community.

## ACKNOWLEDGEMENTS

Figures were created with BioRender.com and Adobe Illustrator.

